# Detecting mosaic patterns in macroevolutionary disparity

**DOI:** 10.1101/423228

**Authors:** Caroline Parins-Fukuchi

## Abstract

Evolutionary biologists have long sought to understand the full complexity in pattern and process that shapes organismal diversity. Although phylogenetic comparative methods are often used to reconstruct complex evolutionary dynamics, they are typically limited to a single phenotypic trait. Extensions that accommodate multiple traits lack the ability to partition multidimensional datasets into a set of mosaic suites of evolutionarily linked characters. I introduce a comparative framework that identifies heterogeneity in evolutionary patterns across large datasets of continuous traits. Using a model of continuous trait evolution based on the differential accumulation of disparity across lineages in a phylogeny, the approach algorithmically partitions traits into a set of character suites that best explains the data, where each suite displays a distinct pattern in phylogenetic morphological disparity. When applied to empirical data, the approach revealed a mosaic pattern predicted by developmental biology. The evolutionary distinctiveness of individual suites can be investigated in more detail, either by fitting conventional comparative models or by directly studying the phylogenetic patterns in disparity recovered during the analysis. This framework can supplement existing comparative approaches by inferring the complex, integrated patterns that shape evolution across the body plan from disparate developmental, morphometric, and environmental sources of phenotypic data.

## Introduction

Characterizing the ways in which phenotypic disparity and evolutionary rates differ across lineages and throughout time has long been a central goal in evolutionary biology. Shifts in the rate of phenotypic change often coincide with the rapid diversification and ecological innovation of diverse lineages (Rabosky and Adams 2012; Rabosky *et al*. 2013). In addition, studying fluctuations in the tempo of evolutionary change can shed light on our knowledge of important evolution processes, such as adaptation and evolutionary constraint. Over the past several decades, the development and application of phylogenetic comparative methods (PCMs) that identify patterns of phenotypic change using phylogenies (Harvey and Pagel 1991; Hansen 1997a; Butler and King 2004; O’Meara *et al*. 2006; Eastman *et al*. 2011; Beaulieu *et al*. 2012). Modern PCMs enable researchers to infer shifts in evolutionary rate, constraint, and disparity throughout time by applying stochastic models of trait evolution to comparative phenotypic data using phylogenetic trees. These approaches have helped to answer important questions concerning the tempo and mode of phenotypic evolution across deep timescales (Harmon *et al*. 2003; Scales *et al*. 2009; Harmon *et al*. 2010; Rohlfs *et al*. 2013; Slater 2013).

Paleobiological and comparative studies have typically examined univariate evolutionary patterns. This has perhaps grown from the tendency of early comparative work towards documenting evolutionary patterns using single key traits, such as molar shape (Simpson 1944; Gingerich 1974). Although such character sampling limits the ability to ask detailed questions relating to morphological evolution, this framework has been used both to explore specific case studies (Scales *et al*. 2009) and broad macroevolutionary questions using single gross morphological traits, often as correlates for overall phenotypic variation (Harmon *et al*. 2003; Beaulieu *et al*. 2007; Harmon *et al*. 2010; Burbrink and Pyron 2010; Rabosky *et al*. 2013; Zanne *et al*. 2014; Bokma *et al*. 2015; Landis and Schraiber 2017).

Although single-trait study models have revealed much about evolution at broad scales, a central goal in post-synthesis evolutionary biology has been to evaluate patterns across anatomical regions and at different phenotypic levels. Such broad investigations require approaches that can accommodate multiple characters. Recent contributions have explored multivariate approaches that enable the analysis of multiple traits simultaneously (Adams 2014a,b,c; Adams and Collyer 2017; Denton and Adams 2015). These enable inference of trait models commonly employed in univariate PCMs among high-dimension data. Another set of methods statistically evaluates correlated evolutionary change between multiple traits along a phylogenetic tree (Revell and Harmon 2008; Caetano and Harmon 2018). These methods will be critical extensions to PCMs moving forward, given the current influx of large, publicly available databases of morphology (Boyer *et al*. 2016), and emerging approaches for the algorithmic and crowd-sourced quantification of variation from digital specimen images (Boyer *et al*. 2015; Chang and Alfaro 2015; Pomidor *et al*. 2016).

One aspect that has been under-explored in a comparative analytical context has been the tendency for distinct subsets of traits to display unique patterns in evolutionary rate and relative disparity. The historical focus on univariate analysis in PCMs and paleobiology may have contributed to a general lack of recognition of the diversity of evolutionary patterns that can combine to organismal body plans. Nevertheless, mosaic evolution has been a fundamental concept in evolutionary biology since the modern synthesis in its acknowledgement of the reality that anatomical regions are often exposed to natural selection at differing magnitudes and directions at different times (Stebbins 1983). The mosaic concept dates to Dollo (Gould 1970), and has frequently been invoked qualitatively as a key factor driving divergent morphological adaptation in different anatomical regions across lineages (De Beer 1954; Cracraft 1970; Mayr 1970; Gould 1977b; Stanley 1979). Although examination in a quantitative context has been limited overall, several key studies have shown that mosaic patterns explain the emergence of important traits in diverse case studies, such as structural variation in the brain across mammals (Barton and Harvey 2000), phenotypic and genomic diversity across angiosperms (Stebbins 1984), and the defining suite of morphological characters displayed by the hominin lineage (McHenry 1975; Gould 1977a; Holloway and Post 1982).

In addition to being of fundamental biological interest, mosaic patterns have long been argued to present unique challenges when inferring phylogeny from morphological characters (Farris 1971). This concern has recently been reasserted by Goloboff and colleagues (2018), who suggest that heterogeneity in relative disparity, as measured by phylogenetic branch lengths, displayed across separate suites of morphological characters confounds phylogenetic inference from morphological traits using Bayesian and maximum likelihood approaches. These concerns parallel the conclusions of important recent studies that validate the prevalence of mosaic patterns in paleobiological and comparative data by manually testing the variability in evolutionary mode that can occur across large morphological datasets (Hopkins and Lidgard 2012; Felice and Goswami 2018). These studies demonstrate the urgent need for overdue extensions to existing uni- and multivariate PCMs that accommodate mosaic patterns in large phenotypic datasets. Methods that separate traits according to overall rate have long been available for phylogenetic inference (Yang 1996; Schraiber *et al*. 2013). However, these approaches do not adequately address mosaic patterns, which focus more on heterogeneity in relative disparity across lineages rather than absolute rate. Despite the importance, there has not yet been a computational approach that algorithmically partitions traits according to their patterns in disparity.

In this paper, I present a novel method that identifies suites of continuous traits displaying shared patterns in disparity to reconstruct the mosaic trends that have shaped organismal phenotypes. The mosaic character suites identified by the approach are defined by partitions of traits that are best explained by shared phylogenetic branch lengths, measured in units of average disparity, along a fixed topology. The phylogenetic models underlying the construction of mosaic character suites, by representing the accumulation of disparity across lineages, thus provide information on relative rates of evolution. After introducing the method, I evaluate its performance using simulated data. I also present an analysis of an empirical dataset of developmental ossification times complied by Rose (2003). This dataset has previously been leveraged to explore mosaic heterogeneity in evolutionary pattern (Germain and Laurin 2009; Laurin 2014) and so is well-suited to an empirical evaluation of the method that I introduce here.

## Methods and Materials

### Code and data availability

The approach described below is implemented in a program called *greedo*. It is available freely on Github at https://github.com/carolinetomo/cophycollapse. All analyses on simulated and empirical data were performed using this program. Scripts and data used for the simulated and empirical analyses are also available on Github (https://github.com/carolinetomo/mosaic).

### Partitioning traits into mosaic suites

The method described here combines several unsupervised learning strategies to partition traits into separate mosaic character suites, with each possessing its own set of phylogenetic branch lengths expressed in units of disparity. These strategies are applied in sequence (Fig. 1), with the goal of identifying the configuration that yields the lowest AIC score.

**Figure 1.**
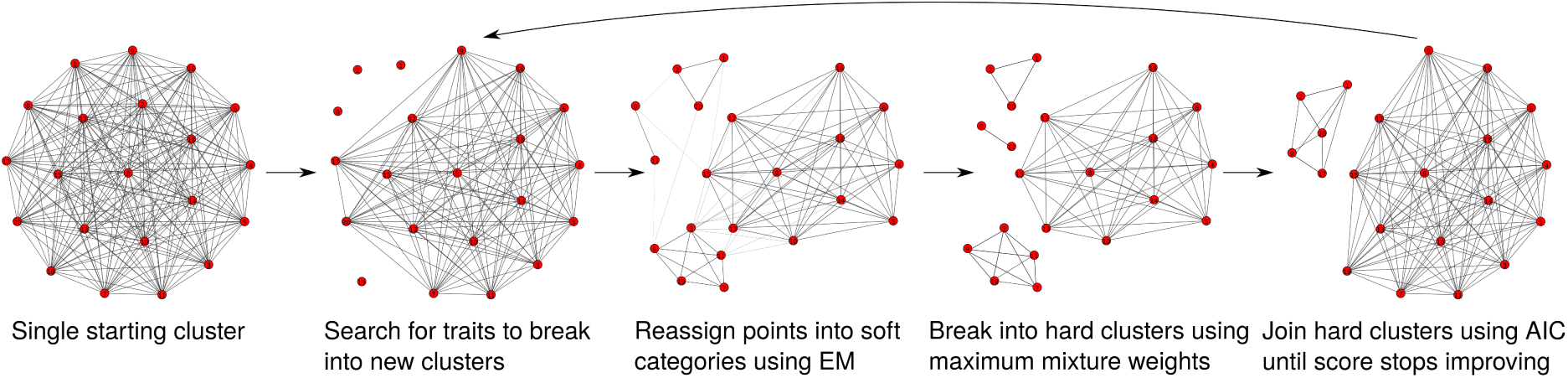
Search procedure to identify mosaic suites.

Each mosaic character suite that defines the classification model contributes to the likelihood independently. The log-likelihoods of each of the traits belonging to character suite *j* are calculated under the associated branch lengths and added to yield an independent log-likelihood. The log-likelihood of the trait matrix, LL_classification_, is calculated by summing the log-likelihoods of all *k* character suites:

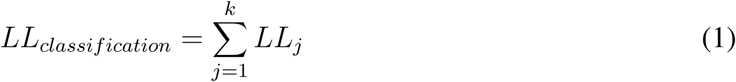

### Tree model

Each mosaic character suite of the classification model is defined by a phylogeny where the topology is fixed, but its branch lengths are free to vary and estimated from the constituent traits. Branch lengths are expressed in units of disparity and are estimated under a Brownian model of trait evolution. The distribution underlying the traits belonging to each partition are assumed to be multivariate Gaussian, with variances between taxa defined by the product of their evolutionary distance measured in absolute time and the instantaneous rate parameter (*σ*^2^). The phylogenetic comparative methods literature often estimates *σ*^2^ alone by assuming a fixed timescale given by branch lengths that have been scaled to absolute time using a clock model (Butler and King 2004; O’Meara *et al*. 2006; Eastman *et al*. 2011). However, here the absolute times are assumed to be unknown, and the rate and time parameters are confounded with one another. Thus, branch lengths are expressed in units of average morphological disparity, or variance per trait.

This parameterization differs from the typical use of stochastic continuous trait models in comparative studies. Traditional approaches generally start by applying single rate (clock-like) Brownian models to chronograms, comparing the fit to more complex models. For instance, *σ*^2^ is sometimes allowed to vary locally in distinct subclades to yield ‘multi-rate’ BM models (O’Meara *et al*. 2006; Eastman *et al*. 2011; Thomas and Freckleton 2012). Expanding upon this, more parameter rich models have also been developed, such as Ornstein-Uhlenbeck (OU) (Hansen 1997b) and jump-diffusion (JD), or Lévy, processes (Landis *et al*. 2012). OU models expand upon standard BM by introducing a term, *α*, that generates a stabilizing force that constrains movement around an optimal trait value, while JD processes contain terms describing the frequency of jumps in character spaces. OU models are often used similarly to multi-rate BM, by allowing optimal trait values to vary across the tree (Butler and King 2004; Beaulieu *et al*. 2012). JD models relax single-rate BM by allowing sudden jumps in mean trait values (Eastman *et al*. 2013; Landis and Schraiber 2017).

The parameterization of the tree and branch lengths used here is similar to that encountered in phylograms reconstructed from molecular and discrete morphological data. In discrete character-spaces, branch lengths are expressed in units of substitutions per site by similarly confounding the rate and time parameters. As used here, this approach has several benefits over the more common comparative approaches described above. Although it may be possible in principle to develop a similar approach that assigns traits to suites associated with separate OU models fit to chronograms with different adaptive regimes, or best explained by distinct BM and JD processes, the tree model used here captures much of the same information (Fig. 2), while simplifying inference from both a statistical and computational standpoint. While multi-rate BM, OU, and JD models are often aimed at describing explicit evolutionary processes, such as environmental adaptation or quantum evolution, they may simply improve model fit over simpler models by accommodating heterogeneity in disparity accumulated across the tree. Nevertheless, more complex models remain useful tools for the exploration of tempo and mode when applied thoughtfully. As a result, the mosaic approach introduced here should be viewed as complementary to existing tools for the examination of tempo and mode. For instance, researchers could delimit mosaic suites of traits using the framework introduced here, and then investigate more detailed patterns by fitting more sophisticated comparative models to the reconstructed suites.

**Figure 2.**
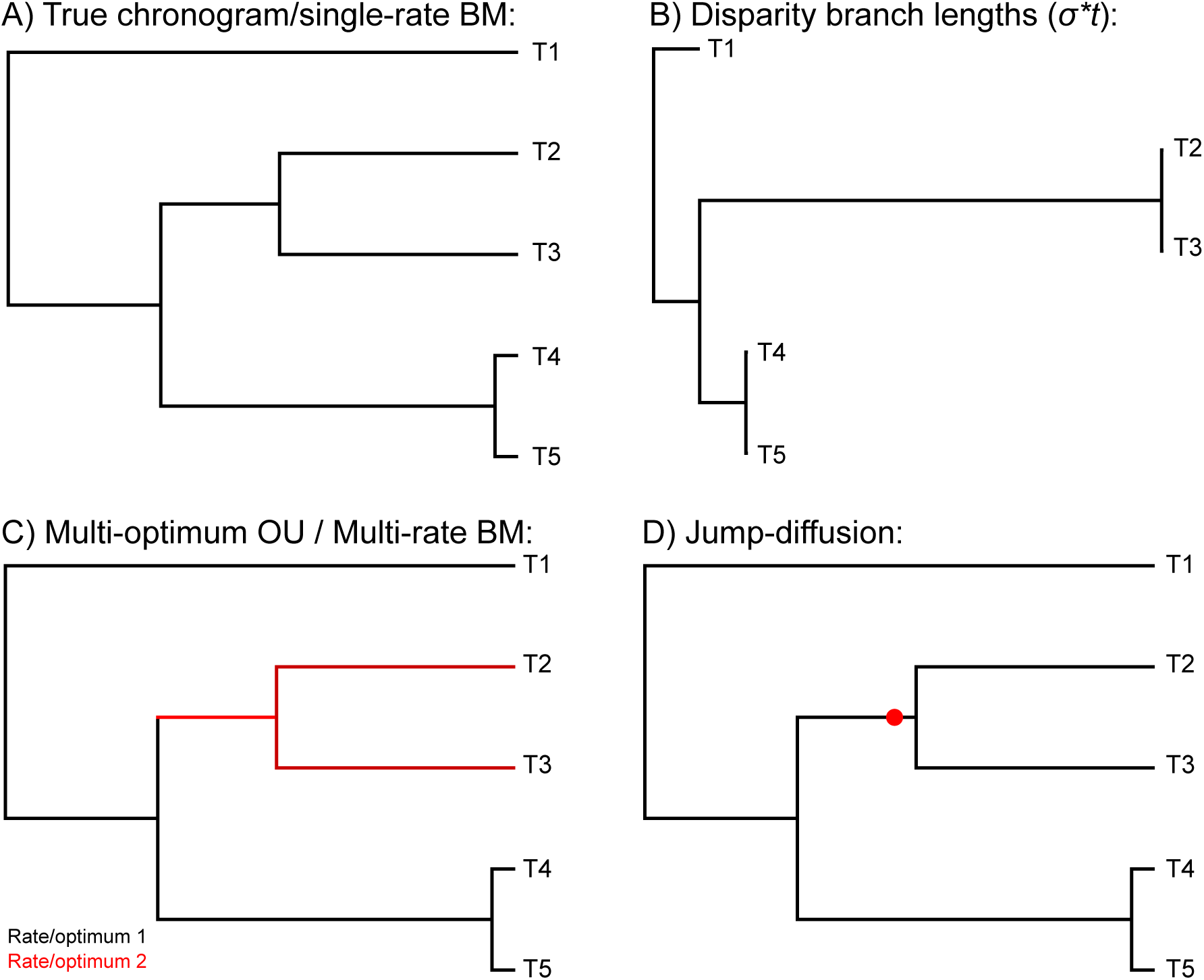
Fitting increased disparity observed in taxa T2 and T3. A) Single rate BM fit to a chronogram assumes clock-like evolution, ignoring both positive jumps in disparity and stasis. B) Expressing branch lengths in units of disparity under the phylogenetic Brownian parameterization used here captures both increased disparity and stasis. C) Fitting multiple Brownian rates or shifts in adaptive optima or D) assuming the presence of a single jump in optimum trait value may explain the same pattern with more model parameters. Note that panels B, C, and D all display similar information, while panel A assumes clock-like evolution.

The likelihood is calculated in a recursion from the tips to the root after Felsenstein (1973). Full derivations of the likelihood and algorithm are also given by Felsenstein (1981) and Freckleton (2012), and summarized briefly here. The tree likelihood is computed from the phylogenetic independent contrasts (PICs) using a ‘pruning’ algorithm. Each internal node is visited in a postorder traversal, and the log-likelihood, *L*_node_ is calculated as univariate Gaussian, with a mean equal to the contrast between the character states, *x*_1_ and *x*_2_ at each subtending edge and variance calculated as the sum of each child edge, *v*_1_ and *v*_2_:

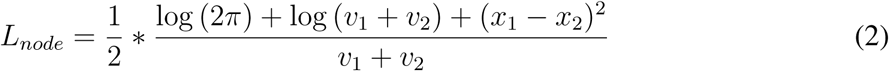

The PIC, *x*_internal_, is calculated at each internal node and used as the character representing the internal node during the likelihood computation at the parent node. The edge length of the internal node, *v*_internal_ is also extended by averaging the lengths of the child nodes.

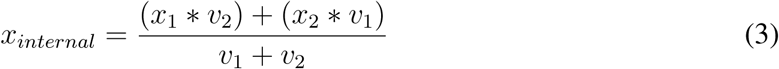

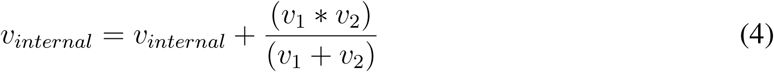

The total log-likelihood of the tree, *L_tree_* is calculated by summing the log-likelihoods calculated at each of the *n* internal nodes.

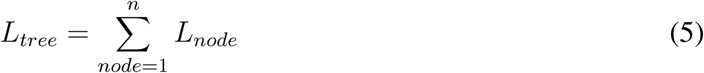

Branch lengths are estimated by iteratively solving the analytical solution to the maximum likelihood (ML) branch lengths for a 3-taxon star topology. In this procedure, the tree is treated as unrooted. Picking a single internal node, PICs are calculated to each of the three connected branches. These are treated as ‘traits’ at the tips of a three-taxon tree. The edge lengths of the pruned tree (*v_i_*) are then computed analytically using the MLE solutions for a three taxon tree (Felsenstein 1981). This procedure is performed on all of the internal nodes. This process is iterated until the branch lengths and the likelihoods converge, yielding a local optimum of the likelihood function. The algorithm and derivation of the 3-taxon ML solutions are given a detailed explanation by Felsenstein (1981).

### Search procedure

All traits start in a single shared partition, where a single model is fit to the entire character matrix simultaneously. From here, traits that display an improved likelihood when evaluated under their own model compared to the model associated with its current cluster are split into their own clusters. A penalty is imposed that is proportional to the difference in size between the existing partitions, as measured in the number of constituent traits. This functions similarly to a Dirichlet process prior by forcing traits to prefer belonging to partitions with a larger number of traits. This constraint is designed to prioritize traits with strongly divergent signal and to discourage overfitting of the clustering model. As a result, only traits with a strong preference for the new component over the existing component are selected. This step is repeated either until the number of occupied categories reaches a user-specified maximum threshold, or there are no more traits left to separate.

From here, the problem is temporarily recast as a finite mixture model, with the number of components corresponding to the user-specified value. First, membership weights are calculated for each trait-component pair as the probability of the trait (x_i_) belonging to each *j* of *K* components. This value is calculated for each component as the proportion of the likelihood of x_i_ (L_ij_) under the corresponding set of branch lengths relative to the summed likelihoods of x_i_ under all *K* components.

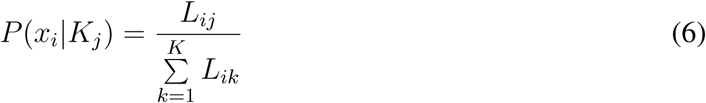

Expectation-maximization (EM) (Dempster *et al*. 1977) is performed to update the mixture weights and the branch length parameters. The entire data matrix is used to estimate the parameters in the phylogenetic model that defines each component, with the contribution of each trait up- or down-weighted according to its probability of membership in each suite (calculated from equation 6). During this step, the model could be thought of as a variation of a typical multivariate Gaussian mixture model, where the covariance matrix is constrained to reflect the structure of a phylogenetic tree, since the phylogenetic Brownian model yields a multivariate Gaussian likelihood function.

Once the mixture model has been updated for several iterations, the components are broken into hard clusters, with the assignment for each trait chosen to be the mixture component that yields the highest likelihood. In the previous step, trait memberships are ‘fuzzy’ in that each trait can make weighted contributions to the phylogenetic models corresponding to multiple clusters. These fuzzy memberships are broken in the current step such that each trait is forced to belong to a single cluster. The clusters are then reduced in an agglomerative manner. At each step of this procedure, the pair of clusters that yields the greatest improvement in AIC, calculated using the classification likelihood (equation 1), is merged. This hierarchical merging continues until either the AIC score cannot be further improved, or only a single component is left. The entire procedure is then repeated from this reduced configuration for a user-specified number of iterations. This number of iterations needed to find the best result is dependent on the specific dataset and contingency of the stochastic rearrangements. Generally, users should feel comfortable truncating the search after the likelihoods cease to improve.

### Stochastically varying mosaic patterns

To examine the strengths and shortcomings of the method in detecting multiple suites of traits that vary stochastically, I performed tests using simulated datasets. A single topology of 20 taxa was simulated under a pure-birth model. I then simulated matrices of continuous traits that each contained multiple partitions of traits displaying distinct simulated disparity patterns. Each partition was simulated under Brownian motion along a tree with distinct branch lengths, generated by drawing randomly from either a gamma or exponential distribution. The rate parameter of the Brownian process was set to 1 across the entire tree so that the matrices reflected the scale and heterogeneity of rates resulting from the altered branch lengths. Each trait matrix contained a single partition simulated under the original ultrametric branch lengths. The randomly drawn branch lengths were intended to mimic the differing rates of evolution that can be experienced by different lineages during evolutionary divergence, with the ultrametric branch lengths reflecting clock-like evolution (Fig. S1). This procedure resulted in highly complex simulated datasets that tested the ability of the method to detect mosaic structure of high dimensionality. All trees and traits were simulated using the phytools package in R (Revell 2012a).

Using this procedure, datasets comprised of 2, 3, and 4 partitions of 50 continuous traits each were generated. All traits were rescaled to a variance of 1. I ran *greedo* on these datasets to attempt to reconstruct these partitions. The maximum number of clusters for these runs was set to half the number of traits in each matrix. This choice of maximum number of clusters should not be interpreted as a prior reflecting the user’s expected number of clusters in the dataset. Instead, this values acts as a mixing parameter meant to place a ceiling on the number of clusters from which to begin the agglomerative merging step of the algorithm. This value can be as high as the number of traits present in the matrix, but can be lowered to speed computation during each iteration. In general, higher values are expected to yield better performance and mixing, but should be chosen to reflect a reasonable trade-off between computation time per iteration and the number of iterations needed to achieve reasonable accuracy.

### Power analysis and detection of relative rate shifts

I performed another simulation experiment to evaluate the statistical power and limits of the method under more controlled conditions. To evaluate the performance of the method computationally, I expanded the size of the simulated trees to 100 taxa. In this trial, I simulated datasets using a procedure modeled after Eastman *et al*. (2011). First, I simulated 100 unique pure-birth trees. I then altered the branch lengths to all be equal, with the trees scaled to a total height of 1. This yielded a set of 100 ‘equal rates’ trees that displayed uniform disparity across all branches. To simulate heterogeneity in evolutionary pattern, I then randomly selected one clade in each of the 100 trees, and multiplied the branch lengths by factors of 8, 16, 32, and 64. Laurin (2014) discovered a 90-fold difference in rate in the most extreme case in his empirical dataset, and so the upper threshold for the shift magnitude chosen here yielded less pronounced heterogeneity than may be expected in real biological systems. As a result, the simulated data represented a reasonable test of the capability of the method to perform well when applied to empirical datasets. The resulting datasets revealed patterns where the clade undergoing the rate shift yielded traits that were mildly (8x) to substantially (64x) differentiated from those generated under the background rate (Fig. S1).

Like in the tests performed by Eastman *et al*. (2011), these rates were inherited across the entire clade. Clades randomly chosen for shifts were constrained to those containing at least 10, and no more than 90, terminal taxa. This resulted in a set of 400 ‘rate shift’ trees of differing magnitude. I then compiled datasets by simulating 50 traits on the trees with equal branch lengths and the trees displaying rate shifts, respectively, with the goal to ascertain the ability of the method to separate traits displaying a shift in rate from those simulated under uniform rates. The 400 resulting datasets contained 100 traits, half of which were simulated along an equal rates tree, and the other half along a rate shift tree of one of the four shift magnitudes. To examine the prevalence of type 1 error, I also simulated datasets of 100 traits along each of the equal branch length trees to test whether the method correctly identified only one suite of traits. Like above, all traits were scaled to a variance of 1 to ensure that the method was correctly identifying heterogeneous patterns in relative disparity, rather than finding heterogeneity reflected in empirical variance, which would unfairly bias results in its favor. This step was also performed in the test described below.

### Distinguishing rate shifts among covarying traits

In nature, many traits are expected to covary. Such covariance has been suggested as a challenge to phylogenetic analyses (Felsenstein 1988) by increasing bias in morphological datasets. To test the behavior of my approach when detecting mosaic patterns among traits that form covarying modules, I performed an additional experiment. I simulated covarying datasets along the 16x and 64x rate shift trees generated above. Again, 50 traits each were simulated along the equal lengths and rate shift trees, respectively, but with the traits separated into two covarying sets of 25 traits. These were compiled into datasets of 100 traits, displaying two distinct evolutionary patterns (equal and shifted rates), and four separate covarying modules. To examine the effect of covariance on type 1 error, I also simulated datasets of 100 traits along the equal rates trees, as above, but separated into four covarying modules. The strength of covariance is likely to affect results, and so data were simulated at correlational intervals of 0.1 (weak), 0.5 (moderate), and 0.9 (strong). Since the approach introduced here does not explicitly model evolutionary or phenotypic integration, this test examined the extent to which these common patterns confound the identification of mosaic patterns in relative disparity. All simulated datasets were generated using the ‘fastBM()’ function in phytools (Revell 2012b).

### Evaluation of reconstruction accuracy

I used the adjusted Rand index (ARI) to evaluate the accuracy of the inferred partitionings (Hubert and Arabie 1985) against the true partitionings. ARI is a corrected version of the Rand index (RI), which measures congruence by counting the pairs of elements that either occupy the same or different clusters in both of the two clusterings, and calculating the proportion of this value relative to all of the possible permutations of elements. As a result, the RI can range from 0, indicating total disagreement, to 1, indicating total agreement. The ARI is similar, but corrects for the potential that for elements to be placed within the same cluster purely due to chance. The ARI usually ranges from 0 to 1, with a value of 0 indicating that a clustering is no more congruent than would be expected from a random assignment of elements to clusters, and 1 indicating complete congruence. The metric takes negative values when a clustering is worse than random. ARI is thus a conservative measure that evaluates the ability for a clustering algorithm to perform more accurately than randomly assigning data points to clusters, rather than the absolute congruence between a reconstructed and true clustering. Nevertheless, it should be noted that random clusterings can frequently display very close congruence to the truth by chance; ARI thus measures the capability of a method to predictably outperform such random sorting. A review of clustering methods found that most standard clustering methods, including K-means, hierarchical clustering, and Gaussian mixture models, most often score between 0 and 0.4 on the ARI (Dahl 2006).

### Comparison to traditional multivariate Gaussian clustering

I also performed analysis on the simulated datasets using the Mclust package (Fraley and Raftery 1999) to explore the capability of standard multivariate clustering approaches in evaluating mosaic macroevolutionary patterns. Mclust is a widely used package that implements several algorithms to learn both the number of clusters and cluster assignments in a model-based framework. Simpler clustering methods have been used previously to delimit modules of continuous traits (Goswami 2006). While *greedo* currently does not explicitly model covariance across traits, but rather clusters based on phylogenetic disparity, Mclust ignores phylogeny while clustering based on covariance across traits. Therefore, the relative performance between the two approaches can illuminate the most effective ways to capture mosaicism in large phenotypic datasets under different scenarios of evolutionary disparity and trait covariance.

### Empirical analysis

To examine the performance of the method on empirical data, I analyzed a dataset comprised of the ossification sequences of 21 cranial bones sampled across 21 taxa obtained from Laurin (2014), initially assembled by Rose (2003). Laurin identified suites of traits displaying distinct phylogenetic patterns using an ‘evolutionary’ principal component analysis (PCA) and also performed a distance-based hierarchical clustering of the data, making these data well-suited to a test of the method introduced here. I was interested in evaluating the ability of my new approach to detect patterns in relative, rather than absolute, rates across lineages, and so I standardized the variance between the traits to 1. The tree presented in the original study was obtained from the authors and used to perform the comparative analyses here.

## Results and Discussion

### Stochastically varying rates

The approach introduced here succeeds at recovering heterogeneous structure in the simulated datasets. Two properties of a clustering analysis are important when measuring the quality of inference: the number of clusters and their membership. When the number of simulated clusters is small, *greedo* typically infers the correct number, with 2 clusters inferred correctly 95% of the time and 3 clusters 77% of the time (Fig 3b). As the number of clusters increases, the method underestimates the number of clusters, with 4 clusters inferred correctly 55% of the time, erroneously reconstructing 3 clusters nearly 45% of the time. An underestimation of the number of clusters may sometimes be preferable, by identifying only categories that are sufficiently differentiated to warrant macroevolutionary investigation. This pattern is seen in other methods aimed at inferring heterogeneous models (Rabosky 2014).

**Figure 3.**
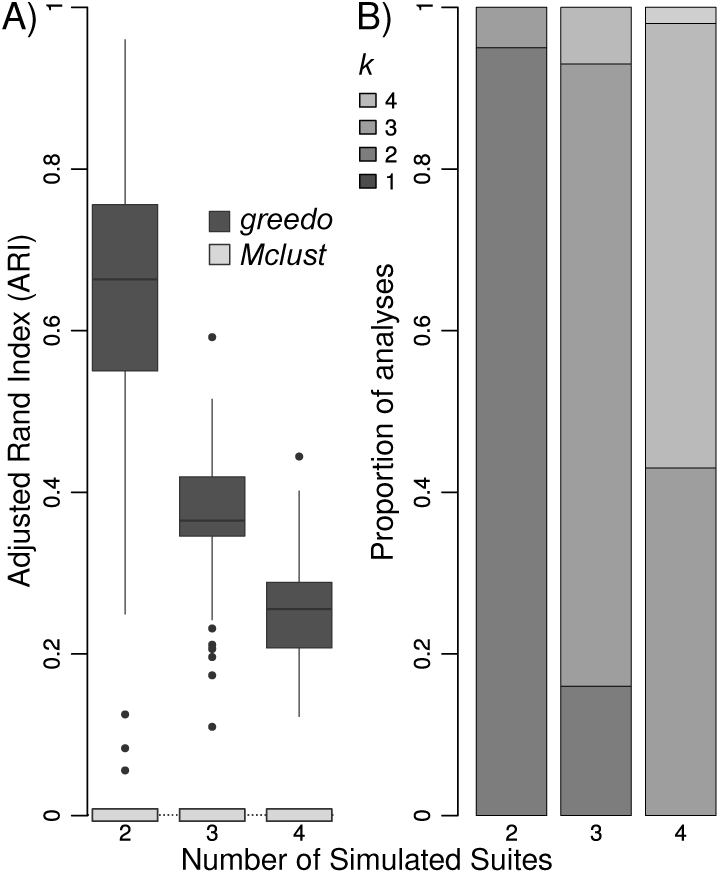
A) Adjusted Rand indices across reconstructions of simulated datasets. Values of zero on the Y axis reflect clusterings that are no closer to the true clustering than ones expected from random assignments. B) Number of clusters resulting from analyses of simulated data. Barplots are stacked to represent the proportion of replicates that resulted in *k* character suites. All barplots are separated by the true number of simulated character suites.

The method also performs well at identifying cluster membership. The two-partition analyses are very accurate, with high ARI values, and nearly always correctly identifying the correct number of clusters. The three- and four-partition analyses were less accurate, but still yield results sufficiently accurate to make macroevolutionary interpretations. ARI values for the three- and four-partition analyses are very similar to published evaluations of general Gaussian clustering approaches, such as Gibbs sampling under a Dirichlet process (Dahl 2006). In analysis of modularity, which are similar in many respects to the patterns targeted here (see below), distance-based hierarchical clustering (Goswami 2006) and PCA (Laurin 2014) have been common tools for machine-guided delimitation of modules. Despite this, I used Mclust for comparison to my new approach. The routine that I used in Mclust models multivariate covariance, and so resembles PCA in the types of patterns that it targets. Mclust was more appropriate for use here because PCA requires human intervention to identify clusters, making it impractical for the analysis of a large number of replicates. Both PCA and Mclust should outperform distance-based clustering approaches, making this a reasonable comparison. Compared to the *greedo* results, Mclust is completely incapable of recognizing heterogeneous structure in this complex dataset. This demonstrates that a phylogenetic approach is needed to identify such subtle, but evolutionarily significant, mosaic patterns.

Despite the encouraging results from the simulated data, the trend toward decreasing accuracy in assignments as components are added suggests either a limitation of the method in adequately exhausting the search space of component assignments or a limitation in the identifiability between clusters as the dimensionality grows. Although the former is a persistent problem in many deterministic clustering algorithms, *greedo*’s use of a stochastic optimization scheme suggests that the latter is likely to be a more pressing challenge as dataset dimensionality grows. As the number of clusters underlying a dataset increases, their defining models will become more similar, eventually approaching a continuous model by compromising the discrete identity of individual clusters. This flattens the optimization surface, weakening the preference of individual traits for their correct cluster (Table 1). As the number of clusters grows, the support for cluster assignment of individual components decreases. In this situation, individual traits may not have enough information to distinguish between membership in separate clusters. This is a common trend in heterogeneous models, where the number of discrete clusters that can inferred is often limited to 4-6 clusters (Yang 1996; Rabosky 2014).

**Table 1.** Median model support in the cluster assignments of each trait resulting from the stochastic rate runs. Support is calculated as the posterior confidence in the recovered suite assignment for each trait.

| k | median confidence |
| --- | --- |
| 2 | 0.9980707 |
| 3 | 0.9539847 |
| 4 | 0.8759063 |

### Rate shifts in independently evolving traits

The method introduced here performs well in distinguishing traits evolved under a single rate shift from those under equal rates (Figs. 4a and 4b). The most subtle rate shift (8x faster) is detectable in 70% of the datasets, although mean accuracy is slightly lower than the 2-suite datasets tested above. This difference is due to decreased power in detecting this more subtle heterogeneity. The method underfits the model in 8x rate shift datasets 30% of the time, by assigning all of the traits to a single shared character suite (Fig. 4b). Power to detect mosaic structure increases with the magnitude of the shift, with the 16x datasets correctly binning the traits into two character suites 95% of the time. The 32x and 64x datasets always correctly identify the number of character suites. Accuracy also increases with the strength of the rate shift. While ARI values in the 8x dataset are all acceptably high, reconstruction error approaches zero as the magnitude of the rate shifts are increased to 64x. Mclust is able to identify some correct structure at very high rate shifts, although it has a tendency to severely overfit the model, inferring 4 or more clusters around 60% of the time in most of the datasets, rarely converging on the correct number of two clusters.

**Figure 4.**
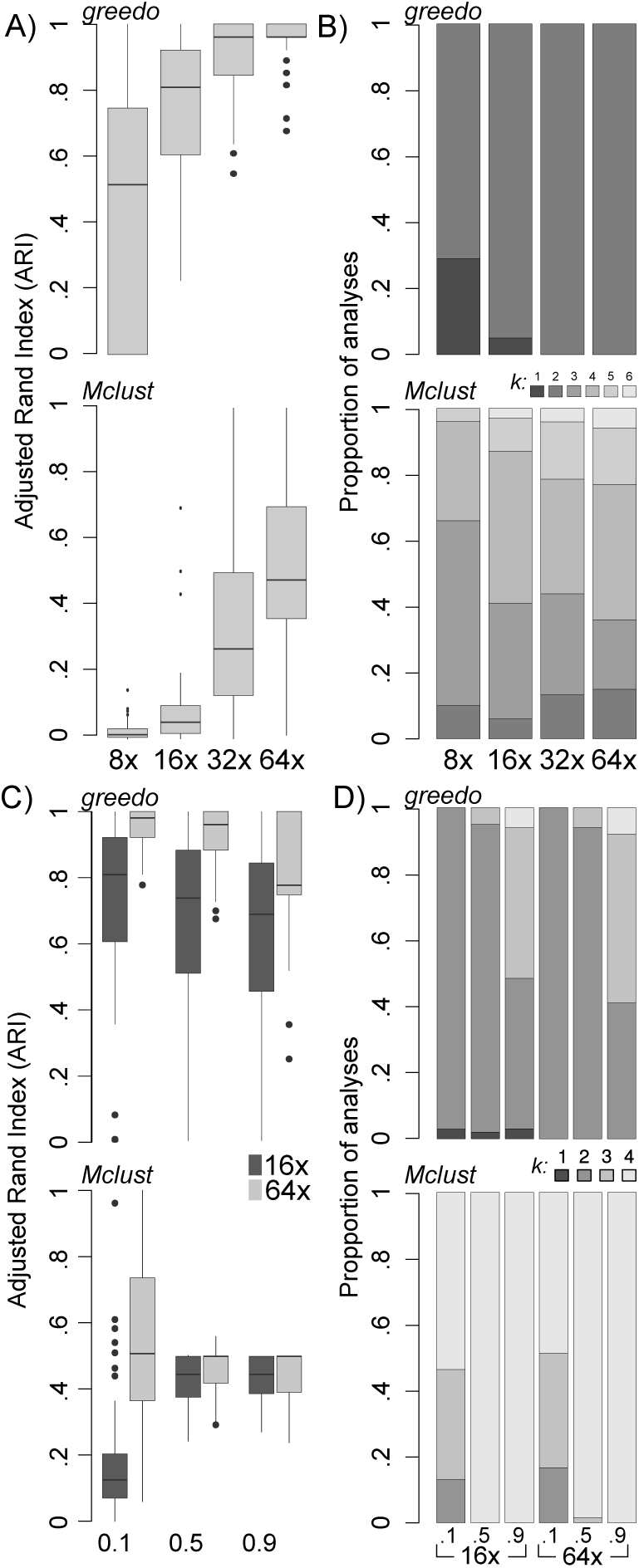
A) Adjusted Rand indices from reconstruction of datasets comprised of mosaic suites with one simulated under equal rates across branches, and the other displaying a randomly placed rate shift at magnitudes of 8, 16, 32, and 64 using both *greedo* and Mclust. B) Stacked barplots to show the frequency that each of *k* clusters are inferred from the simulated datasets. C) and D) show the same metrics as above, summarized from datasets containing simulated covarying modules of characters. The barplots in C are separated by the strength of covariance and colored by the magnitude of the rate shift. All datasets comprised of two mosaic suites.

### Rate shifts in covarying traits

Evolutionary covariance across traits has a detectable, but not overwhelmingly deleterious effect on the capability of the approach in categorizing the number of character suites present in the data (Figs. 4c and 4d). When the simulated modules weakly covary (0.1), the method performs comparably to the independently evolving examples. However, accuracy decreases with the strength of the covariance, with reconstruction being notably less accurate in the strongly covarying datasets. This decrease in accuracy at high levels of covariance can be explained by the ability for *greedo* to detect strongly covarying blocks of characters. While each simulated dataset in this trail was composed of only two macroevolutionary suites, each suite was divided into two blocks of covarying traits. *greedo* can detect covariance at higher intensities, causing the approach to overfit the datasets by inferring the presence of more than two character suites in 59% of cases (Fig. 4d). Mclust performs very well at identifying the covarying modules of traits at moderate to high levels of covariance (0.5 and 0.9). The poor performance at low covariance (0.1) and subtle evolutionary heterogeneity (16x) suggests that, while Mclust performs very well in its intended function, standard clustering approaches are inadequate for the study of mosaic evolutionary patterns. The apparent capability of *greedo* to identify patterns in both covariance and phylogenetic disparity stands in contrast to the poor performance of Mclust under low to zero covariance and suggests that phylogenetic approaches should be preferred. Put simply, although empirical applications may extend both *greedo* and standard clustering approaches outside of the range of functions intended by their underlying models in different ways, *greedo* performs better at identifying a more diverse array of patterns.

### Type 1 error

The method correctly identifies the lack of heterogeneity when no or little covariance is present. As with the multi-rate covarying examples above, the method has a tendency to overfit datasets that display high levels of covariance. It should also be noted that the extremely high and rampant levels of covariance that were needed to induce overfitting in both the type 1 and type 2 tests may be unlikely in many types of empirical data.

### Mosaic evolution in urodele cranial development

Understanding the mosaic evolution of skeletal development has long been a goal in comparative zoology (Goswami *et al*. 2009; Laurin 2014). Application of my new approach to the urodele cranial ossification data demonstrate its potential to contribute to an improved understanding of the mosaic nature of developmental evolution. Four separate runs each yielded different partitionings into two mosaic suites. All arrangements overlap in their assignments, and the AIC scores are all close to one another. To visualize the overall support for the categorization of each trait across partitionings, I calculated the AIC weight of each model (Burnham and Anderson 2002). The AIC weight of model *i*, w_i_ can be interpreted as its probability of being the best model among a set of *K* candidates.

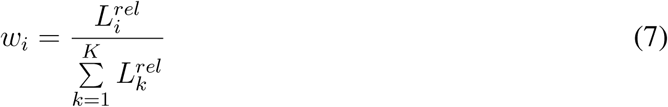

where L_i_^rel^ is the relative likelihood of model *i*:

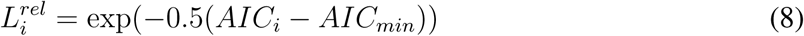

These weights were used to visualize the strengths of the support connecting traits across all the four best partitionings in a graph (Fig. S2). An edge was drawn between traits *i* and *j* if they occurred in the same component in any of the four results, with a weight given by the summed AIC weights of all of the configurations where *i* and *j* occur in the same character suite. The maximum weight possible is 1.0, when traits *i* and *j* share a suite in all of the configurations. The resulting graph suggests that traits 0, 1, 2, 17, and 20 all form a suite, with the rest of the traits sharing a separate suite. This result is very close to the pattern reconstructed by Laurin (2014) using an ‘evolutionary PCA’ approach, differing only in the assignment of the stapes (Table 2). This is likely because the approach developed by Laurin seeks to join traits that display similar values in phylogenetic independent contrasts (PICs), and so focuses on patterns similar to the phylogenetic Brownian likelihood model used here.

**Table 2.** Character suite assignments from original study (Laurin 2014) and the weighted graph in Fig. 3e. Suites are given arbitrary labels that indicate co-occurrence of traits in each study. The two arrangements differ only in the assignment of the stapes.

| Index | Bone | Laurin 2014 suite | *greedo* suite |
| --- | --- | --- | --- |
| 0 | coronoid | 1 | 1 |
| 1 | vomer | 1 | 1 |
| 2 | palatine | 1 | 1 |
| 3 | dentary | 0 | 0 |
| 4 | premaxillary | 0 | 0 |
| 5 | prearticular | 0 | 0 |
| 6 | squamosal | 0 | 0 |
| 7 | parasphenoid | 0 | 0 |
| 8 | frontal | 0 | 0 |
| 9 | parietal | 0 | 0 |
| 10 | pterygoid | 0 | 0 |
| 11 | exoccipital | 0 | 0 |
| 12 | maxilla | 0 | 0 |
| 13 | quadrate | 0 | 0 |
| 14 | opisthotic | 0 | 0 |
| 15 | prefrontal | 0 | 0 |
| 16 | prootic | 0 | 0 |
| 17 | stapes | 0 | 1 |
| 18 | orbitosphenoid | 0 | 0 |
| 19 | nasal | 0 | 0 |
| 20 | septomaxilla | 1 | 1 |

In his original study, Laurin (2014) also performed an exploratory hierarchical clustering of the ossification times and found substantial differences in structure as compared to that revealed by his evolutionary PCA approach. The results here differ from Laurin’s exploratory hierarchical clustering, instead aligning very closely to the evolutionary PCA approach. This is reassuring for the performance of *greedo*, as Laurin considered the evolutionary PCA to yield the correct answer and the hierarchical clustering to demonstrate the inadequacy of raw similarity in delimiting meaningful patterns (Laurin, pers. comm.).

The sole difference between Laurin’s and my results are in the assignment of the stapes. The placement of the stapes in a suite along with the coronoid, vomer, and palatine in my analysis suggests that these four bones share a distinct macroevolutionary pattern from the remaining bones (Fig. 6). Despite the shared pattern in disparity reconstructed here, Laurin’s result suggests that the stapes likely does not covary with the other three bones. Laurin’s evolutionary PCA, which relies on hypothesis testing to identify statistically significant covarying modules, therefore places the stapes in the larger, more inclusive suite. The difference therefore likely stems from *greedo*’s ability to reconstruct shared disparity patterns that result from both statistical covariance and other processes that may drive shared patterns in evolutionary rate in traits that are ontogenetically independent.

**Figure 5.**
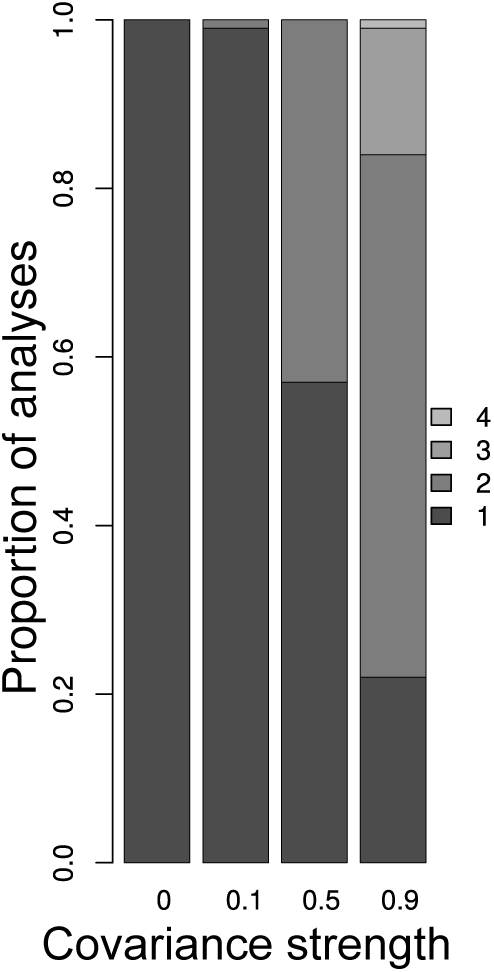
Type 1 error. Reconstructions show no type 1 error when traits are evolved independently or with weak covariance. Type 1 error increases at moderate and high levels of covariance, as the method overfits biased patterns to place covarying modules in superfluous mosaic suites.

**Figure 6.**
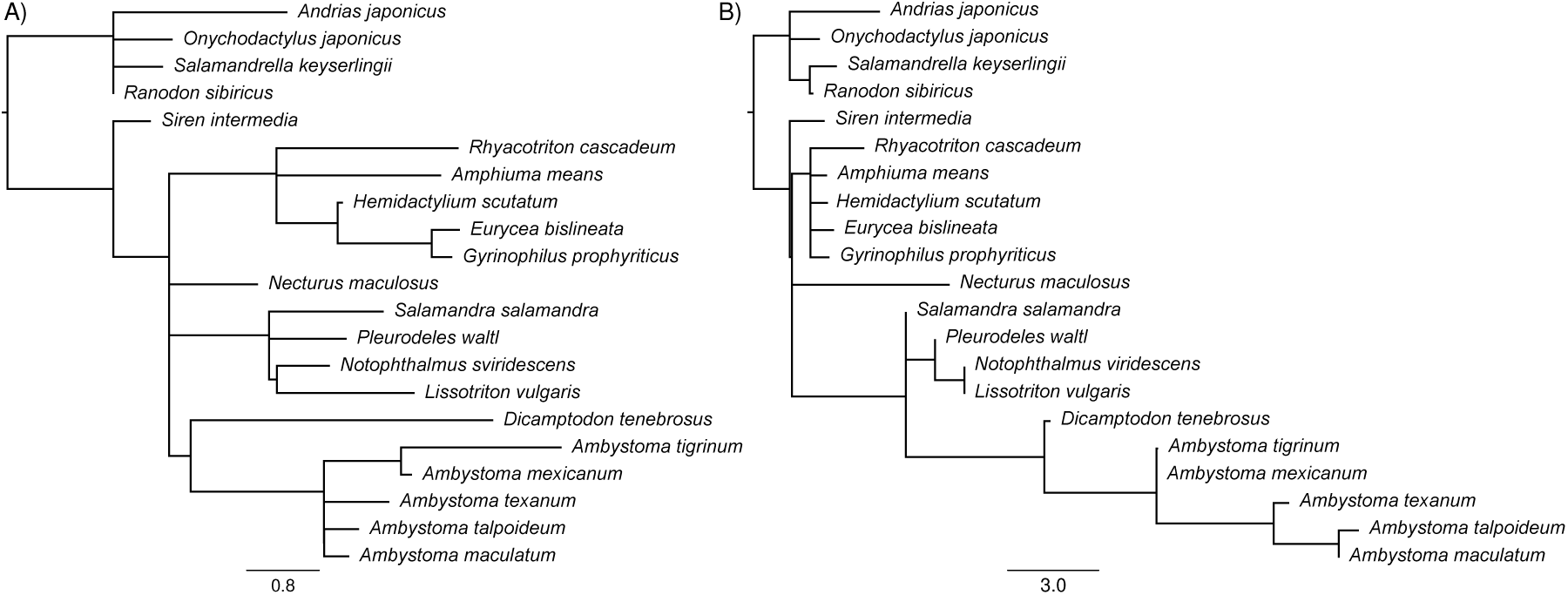
Branch lengths reconstructed from traits contained within: A) suite 0 (Table 2); B) suite 1 (Table 2) display distinct patterns in disparity.

The composition of the mosaic suites recovered by *greedo* shed light on the widely divergent patterns in developmental evolution across cranial bones. By estimating the phylogeny-corrected variance of each bone, Laurin discovered that the stapes displayed a 90-fold increase in absolute evolutionary rate in comparison to the dentary. Here, these two traits occupy separate suites. Since I rescaled the traits to unit variance, the accumulation of disparity across lineages, expressed as branch lengths (Fig. 6), reflects relative, rather than absolute disparity. Despite the difference in scaling and parameterization (disparity versus rates) between my and Laurin’s analysis, both yield similar results. In my analysis, the cluster occupied by the palatine displays a substantially higher accumulation of evolutionary disparity overall, as measured by tree depth, compared to the cluster occupied by the dentary (Fig. 6). The substantial differences in evolutionary signal displayed by the two suites suggests that the evolution of the urodele skull was shaped in part by the substantial evolutionary developmental flexibility displayed by a subset of smaller bones. At the same time, a separate suite of bones, many of which are involved in important ecological functions (for example, mastication in the case of the dentary), remained more conservative in both overall rate and relative divergence across lineages. These distinct patterns in developmental lability and constraint are likely important to the diversification of ecomorphological function in the urodeles. Comparison of this developmental result to those obtained from ecological and adult skeletal data will contribute to a more comprehensive understanding of how the mosaic patterns in developmental evolution revealed here can shed light on the emergence of novelty in vertebrate body plans and ecomorphology.

Although this work provides only a cursory view of the mosaic structure underlying the macroevolution of the cranium, the results are encouraging. As was noted by Laurin, developmental modules have been notoriously difficult to delimit, demonstrating the power of the framework introduced here to illuminate the mosaic origin of variation in a diverse array of phenotypic data. Moving forward, it will be interesting to learn whether shape and size variables representing each of the bones in the developmental sample analyzed here 1) conglomerate to form suites similar in composition and 2) display macroevolutionary patterns in disparity that are similar or distinct to those reconstructed in Figure 7. This will help to determine whether evolutionary changes in developmental suites correspond to innovation in phenotypic suites. Generally, digging deeply into the mosaic complexity underlying the parallel patterns in skeletal development and variation will contribute to a stronger understanding of the interplay between development and functional morphology in shaping organismal diversity.

### Utility to phylogenetic comparative methods

Previous work has shown that morphological (Lynch 1990) traits often display patterns in rate that are not easily distinguishable from conservative evolutionary forces such as genetic drift and stabilizing selection. However, mosaic evolution suggests that such conservatism may vary throughout time and across the body plan. As an exploratory method that delimits suites of traits that are functionally and/or developmentally integrated, the approach that I introduce here can identify the most important axes of evolutionary variation across many characters when applied to morphometric and developmental data. More complex models can then be fit separately to the resulting mosaic suites to explore how distinct evolutionary processes have acted at different times in separate suites. Simultaneous application of this framework to morphometric and developmental data will spur a deeper understanding of the origins of phenotypic novelty by characterizing the conditions under which specific integrated suites tend toward evolutionary innovation. For example, comparison of the mosaic patterns displayed by skeletal measurements to those displayed by ossification data may help to understand whether rapid bursts in phenotypic differentiation correspond to concomitant shifts in the evolution of development.

### Utility for phylogenetic inference and divergence time estimation

In addition to being biologically interesting in their own right, the mosaic suites identified by my approach will be useful in methods for inferring phylogeny and divergence times. Several recent articles have demonstrated the strong potential for continuous traits as an alternative to discrete traits when reconstructing phylogeny (Parins-Fukuchi 2018b,a) and divergence times (Alvarez-Carretero *et al*. 2018). Despite these advances, one persistent issue in morphological phylogenetics has been the difficulty in accommodating the expected heterogeneity in branch lengths across traits. Cladists have argued this to be a fundamental limitation to model-based approaches when demonstrating the prevalence of these patterns across empirical datasets (Goloboff *et al*. 2018). Error stemming from the effect of mosaic branch lengths may be expected to exhibit an even greater effect on divergence time estimation, which depends on calculating rates from branch-wise patterns in disparity.

Although long postulated to be a problem in phylogenetic inference, there has not yet been a computational approach that delimits suites of traits displaying shared patterns in evolutionary rates across lineages. If incorporated into existing methods for phylogenetic reconstruction and divergence time estimation, the approach introduced here would help to alleviate this source of error. Identifying suites of traits based on patterns in relative evolutionary rate may also provide a framework through which to separate clock-like characters from those displaying more erratic patters. Filtering character data based on conformity to clock-like patterns improves divergence time estimation in molecular data (Smith *et al*. 2018), and so would also be beneficial in morphological data. Finally, although the implementation here analyzes patterns in continuous traits, an extension to discrete traits would be straightforward. This would broaden the applicability of the approach to discrete morphological characters and molecular sequence data.

### Covariance, mosaic evolution, and modularity

The approach introduced here does not explicitly model covariance among characters. However, when data are simulated as sets of strongly covarying modules, the method splits the covarying modules of traits evolved along the same set of branch lengths into multiple character suites, rather than combining them. Although this may be seen as overfitting in some respects, this behavior may be useful in practice. While standard approaches such as Mclust and PCA (both phylogenetic and non-phylogenetic) focus on covariance alone, *greedo* is capable of identifying both mosaic patterns as reflected in branch lengths and covarying suites of traits. This is an encouraging result, since the mosaic patterns in branch lengths explicitly modelled by the method may be seen as a superset of patterns that stem from covariance alone. Ideally, future extensions to this method will explicitly model covariance by incorporating older or recent approaches to identifying evolutionary covariance across multiple continuous traits (Felsenstein 1973). Nevertheless, even in its current form, the analyses here show that *greedo* will be a useful alternative to traditional covariance-based approaches through its ability to place both covarying and independent traits into a macroevolutionary context during the initial exploratory steps when analyzing large datasets of traits.

Although Mclust performs well when applied to datasets that display high covariance, it fails at recognizing heterogeneous structure in the presence of low to zero covariance. Standard clustering approaches have previously been applied to the delimitation of modules in morphological data (Goswami 2006), but the poor performance of Mclust overall echoes the finding of Laurin (2014) that traditional clustering is inappropriate for analyses of mosaicism and modularity. Through its ability to identify both covarying and shared macroevolutionary structure, the general framework introduced here is better suited to the identification of mosaic macroevolutionary patterns across empirical datasets that may display complex patterns in both disparity and covariance than traditional statistical approaches. Laurin (2014) developed similar arguments, proposing the use of phylogenetic approaches, such as phylogenetic independent contrasts (PICs) Felsenstein (1985), over standard clustering and PCA.

Although similar in spirit, the approach introduced here advances modular analysis of phenotype over PIC-based approaches in several major ways. PICs assume that characters evolve according to single-rate BM. The resulting variances in the PICs can be used to compare rates across different traits, but are less useful for the comparison of rates across taxa. *greedo* allows evolutionary rates to vary across the tree (Fig. 2), allowing for finer resolution into the patterns in trait evolution that define each suite. In addition, phylogenetic PCA and PIC-based approaches are poorly equipped to handle datasets with missing data. The algorithm used here is designed to accommodate missing traits through the inclusion of a EM-based data imputation step when estimating the branch lengths associated with each cluster. This makes the approach well-poised to handle paleontological datasets, which frequently have many missing values. Finally, unlike standard covariance-based approaches (such as PCA), the framework here can be easily extended to discrete characters, widely expanding the range of possible applications to paleontological and neontological questions across the tree of life.

The method introduced here is designed to identify shared mosaic patterns in evolutionary disparity. This aim is closely related to, but subtly distinct from, the existing body of work that aims to identify correlational patterns in morphological integration and modularity (Olson and Miller 1958; Cheverud 1982; Goswami 2006; Goswami *et al*. 2009; Denton and Adams 2015; Adams 2016). The mosaic suites identified here are a superset of the modules identified in typical morphological integration studies. Strong covariance between traits implies the presence of a shared underlying pattern of macroevolutionary disparity (Felice *et al*. 2018). However, as is demonstrated by the simulation experiment performed here, independent modules of covarying traits may share macroevolutionary patterns. While the explicit accommodation of covariance should be a long-term goal for my method, the surprising capability of the method to identify covarying modules of traits as-is suggests its utility in evaluating evolutionary mosaicism in data such as geometric morphometric landmarks, which often exhibit high levels of integration and covariance. This behavior suggests that the framework introduced here will be useful for analyses of modularity by revealing patterns that stem from both covariance and shared evolutionary rates.

The incapability of existing methods to explore the latter has been invoked of a major criticism of existing approaches to modularity (Laurin 2014). By simultaneously identifying structure resulting from both shared rates and covariance, *greedo* can be harnessed as an initial exploratory step. It would then be possible to reconstruct the covariance structure within the resulting mosaic suites to explore more detailed patterns in integration and allometry. It will also be valuable in the analysis of traits quantified using standard linear morphometric approaches and developmental data, such as the ossification dataset explored here. In the future, the method will be useful as applied to data generated using emerging methods that automatically quantify shape differences in specimen images (Pomidor *et al*. 2016).

## Conclusions

While there have been fundamental and seminal works sketching out major historical trends in morphological disparification, most offer only a glimpse into the diversity of patterns that shape separate anatomical regions and operate at different phenotypic levels that extend from the gene to the environment. Although a foundational concept in post-synthesis evolutionary biology, researchers have been limited in their ability to explore mosaic evolution by both the difficulty in assembling comprehensive phenotypic datasets, and the lack of computational methods to handle comparative problems of significant breadth. Emerging approaches that can quantify morphology across entire organisms along with the increasing availability of large-scale transcriptomic and environmental datasets offer the unprecedented opportunity to develop a synthetic evolutionary picture that includes complex mosaic patterns operating at separate phenotypic levels and timescales. Methods like the one introduced here will be critical to these developments by facilitating the reconstruction of the diverse mosaic patterns that have shaped evolutionary variation at different phenotypic levels across the tree of life.

## Acknowledgements

I greatly thank M Laurin, J Saulsbury, SD Smith, SA Smith, N Walker-Hale, Associate Editor DC Collar, and three anonymous reviewers for comments that greatly improved the manuscript.

## Supplementary information

**Figure S1.**
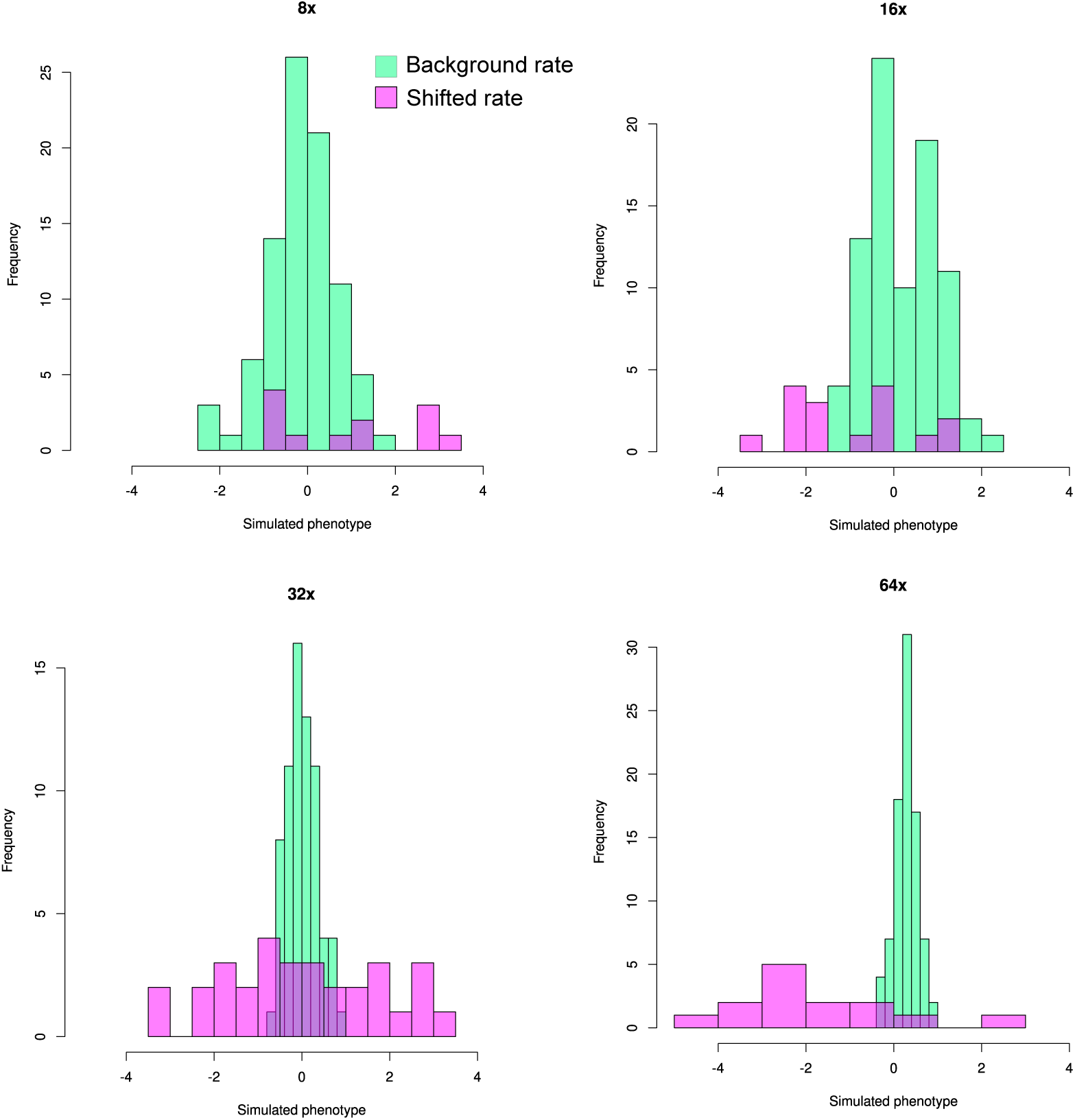
Example of the distributions of traits generated under the shifted and background rate regimes.

**Figure S2.**
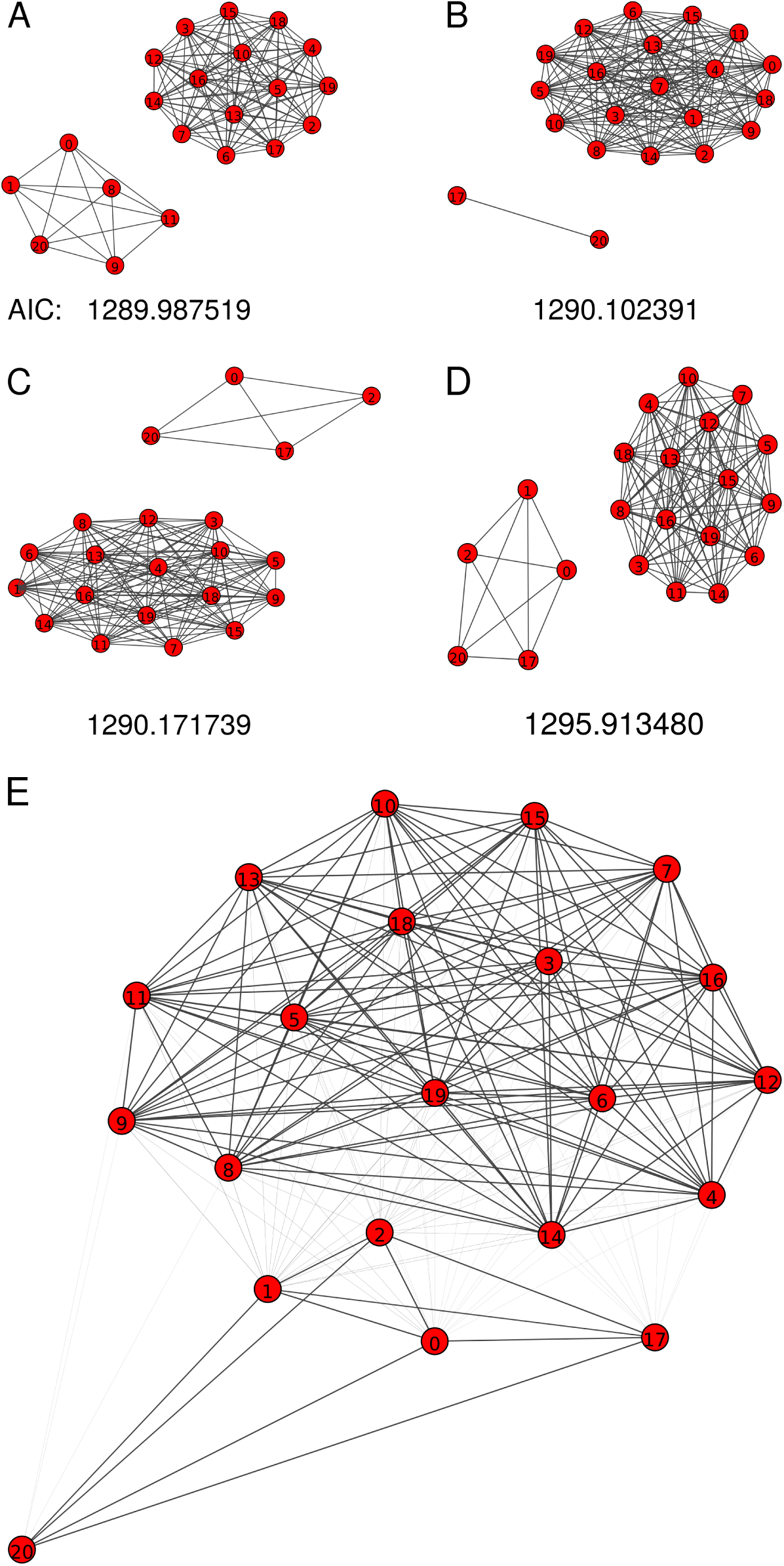
A-D) Four best configurations with AIC scores. E) Weighted graph calculated by summing the AIC weights associated with the each model to form edges and edge weights. Trait numberings correspond to those given in table 2. All graphs were drawn using the “lgl” format implemented in igraph.

### Scale and rate

The approach described here seeks to identify suites of traits sharing similar patterns in relative evolutionary disparity across lineages. Continuous traits displaying greater empirical variances will display higher absolute rates of evolution when modeled under Brownian motion. As a result, I normalized the variances of the datasets used in the simulation experiments and empirical case study. Nevertheless, alteration of the scale of continuous traits may often change the interpretation of results, and so should be performed thoughtfully. In cases where phenotypes are quantified using a single, shared set of units, such as with geometric morphometric data, standardization of the variances across traits erases information characterizing absolute evolutionary rate. In such carefully constructed datasets, including the matrix of developmental sequences used in the empirical example above, researchers may wish to quantify differences in absolute evolutionary rate across characters. For instance, using the same dataset, Germain and Laurin (2009) demonstrated substantial variability in absolute rate across traits. Study of absolute and relative rates can each yield unique insights into evolutionary processes, and so the scaling of traits should be considered carefully. I did not explore inference of heterogeneity in absolute rates here because it is an easier clustering problem, and thus would have been an overly favorable test of the method that I introduced. In addition, approaches that identify heterogeneity in absolute rate have been well explored in the molecular phylogenetics literature (Yang 1996), and successfully ported to continuous traits (Schraiber *et al*. 2013).

### Information criteria and overfitting

In the analyses performed here, I exclusively used the AIC, in lieu of the corrected version, AICc, and the Bayesian Information Criterion (BIC). Previous authors have suggested that the AICc should be generally preferred to the uncorrected version (Burnham and Anderson 2002). My preference for the AIC was driven by several factors. The number of clusters is generally completely unknown prior to the analysis, and perhaps more importantly, there is generally no single ‘true’ clustering underlying the mosaic evolutionary patterns sought by the method. As a result, it might generally be preferable in the context of addressing comparative questions to identify a small number of spurious components in the final configuration than to ignore important biological variation that could be missed due to the steeper penalty imposed by the AICc. The analyses here support this justification. The simulated analyses show that, when AIC is used, overestimating the number of components is not a major problem (Fig. 2). In addition, the results of the empirical analysis suggest that more coherent patterns emerge when several well-supported configurations are averaged. If spurious partitions are encountered in some arrangements, averaging over the results should generally reveal reasonably strong connections between points occupying overfit components.

Although BIC has been used successfully to select the number of components in mixture models (Fraley and Raftery 1998, 1999), I preferred the behavior and basis of AIC for these analyses. BIC assumes that the true model is within the set of candidate models, and so can be sensitive to model-misspecification (Wagenmakers and Farrell 2004). This assumption is incompatible with the goals of my method, which does not seek to identify a single ‘true’ configuration, but instead characterize the major axes of heterogeneity in disparity across lineages. This goal is more consistent with AIC, which simply seeks to identify the model that yields the lowest amount of information loss relative to the dataset. Despite my preference for AIC in the analyses presented here, AICc or BIC may be more appropriate in other situations. As such, researchers should be thoughtful in their choice of information criterion when performing the approach introduced here.

